# Viral origin of eukaryotic type IIA DNA topoisomerases

**DOI:** 10.1101/2022.04.12.488027

**Authors:** Julien Guglielmini, Morgan Gaia, Violette Da Cunha, Alexis Criscuolo, Mart Krupovic, Patrick Forterre

## Abstract

Type II DNA topoisomerases of the family A (Topo IIA) are present in all bacteria (DNA gyrase) and eukaryotes. In eukaryotes, they play a major role in transcription, DNA replication, chromosome segregation and modulation of chromosome architecture. The origin of eukaryotic Topo IIA remains mysterious since they are very divergent from their bacterial homologues and have no orthologues in Archaea. Interestingly, eukaryotic Topo IIA have close homologues in viruses of the phylum *Nucleocytoviricota*, an expansive assemblage of large and giant viruses formerly known as the nucleocytoplasmic large DNA viruses (NCLDV). Topo IIA are also encoded by some bacterioviruses of the class *Caudoviricetes* (tailed bacteriophages). To elucidate the origin of the eukaryotic Topo IIA, we performed in-depth phylogenetic analyses combining viral and cellular Topo IIA homologs. Topo IIA encoded by bacteria and eukaryotes form two monophyletic groups nested within Topo IIA encoded by *Caudoviricetes* and *Nucleocytoviricota*, respectively. Importantly, *Nucleocytoviricota* remained well separated from eukaryotes after removing both bacteria and *Caudoviricetes* from the dataset, indicating that the separation of *Nucleocytoviricota* and eukaryotes is probably not due to long branch attraction artefact. The topology of our tree suggests that the eukaryotic Topo IIA was probably acquired from an ancestral member of the *Nucleocytoviricota* of the class *Megaviricetes*, before the emergence of the last eukaryotic common ancestor (LECA). This result further highlights a key role of these viruses in eukaryogenesis and suggests that early proto-eukaryotes used a Topo IIB instead of a Topo IIA for solving their DNA topological problems.

## Introduction

DNA topoisomerases are ubiquitous enzymes that are essential for solving topological problems inherent to the helical structure of DNA (Champoux 2001; Forterre and Gadelle 2009; Wang 2009; Vos et al. 2011; Forterre 2012; Pommier et al. 2016). Based on mechanistic properties, they have been classified into types I and II. Type I DNA topoisomerases (Topo I) produce transient single-strand breaks in double-stranded DNA (dsDNA) and catalyze the transfer of one DNA strand through this break. In contrast, type II DNA topoisomerases (Topo II) produce transient double-strand breaks and catalyze the transfer of a dsDNA segment (either from the same or different dsDNA molecule) through this break. Five different families of DNA topoisomerases have been defined based on amino-acid sequences and structural similarities: three families of Topo I (Topo IA, Topo IB and Topo IC)(Champoux 2001; Forterre 2006), and two families of Topo II (Topo IIA and Topo IIB) (Bergerat et al. 1997; Gadelle et al. 2003). All Topo II and some Topo I (IB and IC) can relax positive and negative superturns that otherwise would accumulate in front and behind the replication forks and transcription bubbles, respectively. In addition, Topo II can eliminate the catenanes that can accumulate at the end of the chromosome replication. In eukaryotes, Topo IIA are also intrinsic structural components of the chromosomal scaffold (Hizume et al. 2007) and play a major role in modulating chromosome architecture and long-range chromatin structure (Nitiss 2009; Nielsen et al. 2020).

DNA topoisomerases have been extensively studied because they are the targets of important antibiotics and antitumor drugs (Pommier 2013). These drugs interfere with the breakage-reunion mechanisms of the enzyme and transform the transient intermediate topoisomerase-DNA covalent complexes into stable poisons, interfering with replication and transcription. However, these enzymes are also very interesting (and intriguing) in terms of the history of life on our planet. Indeed, the distribution patterns of the different DNA topoisomerase families within the three domains of life, Archaea, Bacteria and Eukarya (eukaryotes), do not fit the usual distribution of informational proteins, such as ribosomal proteins or DNA-dependent RNA polymerases (Da Cunha et al. 2017), raising challenging questions and prompting unorthodox hypotheses (Forterre and Gadelle 2009). Whereas informational proteins from eukaryotes usually much more closely resemble their archaeal homologs than their bacterial ones, the universal eukaryotic Topo II (member of the Topo IIA family) has no obvious orthologue in Archaea. All archaea (except for certain Thermoplasmatales) contain an enzyme of the Topo IIB family, dubbed DNA topoisomerase VI (Topo VI), suggesting that the Last Archaeal Common Ancestor (LACA) contained no Topo IIA but a Topo IIB for relaxation of positive superturns and chromosome decatenation (Forterre and Gadelle 2009). All bacteria encode a unique Topo IIA, DNA gyrase, which is a distant homologue of the eukaryotic enzyme. DNA gyrases are heterotetramers composed of two subunits (GyrA and GyrB) that are homologous to the C-terminal and N-terminal moieties of the homodimeric Topo IIA of eukaryotes, respectively (Fig. 1). Some Archaea encode a two-subunit DNA gyrase very similar to the bacterial enzyme and highly divergent from the eukaryotic Topo IIA. Phylogenetic analysis has indicated that these DNA gyrases were recruited from Bacteria by lateral gene transfer (Forterre et al. 2014). Similarly, some eukaryotes, such as *Archaeplastida*, encode a bacterial-like DNA gyrase (Topo IIA) present in chloroplasts and mitochondria that was most likely acquired from *Cyanobacteria* during the endosymbiotic event that led to the emergence of the chloroplasts (Wall et al. 2004). These eukaryotic Topo II are very similar to their bacterial counterparts and in phylogenetic analyses are nested within the clade of bacterial gyrases (Forterre et al. 2007). It seems unlikely that the very divergent eukaryotic Topo IIA originated through a similar endosymbiotic pathway. The origin of the Topo IIA in eukaryotes thus remains enigmatic.

**Fig. 1.**
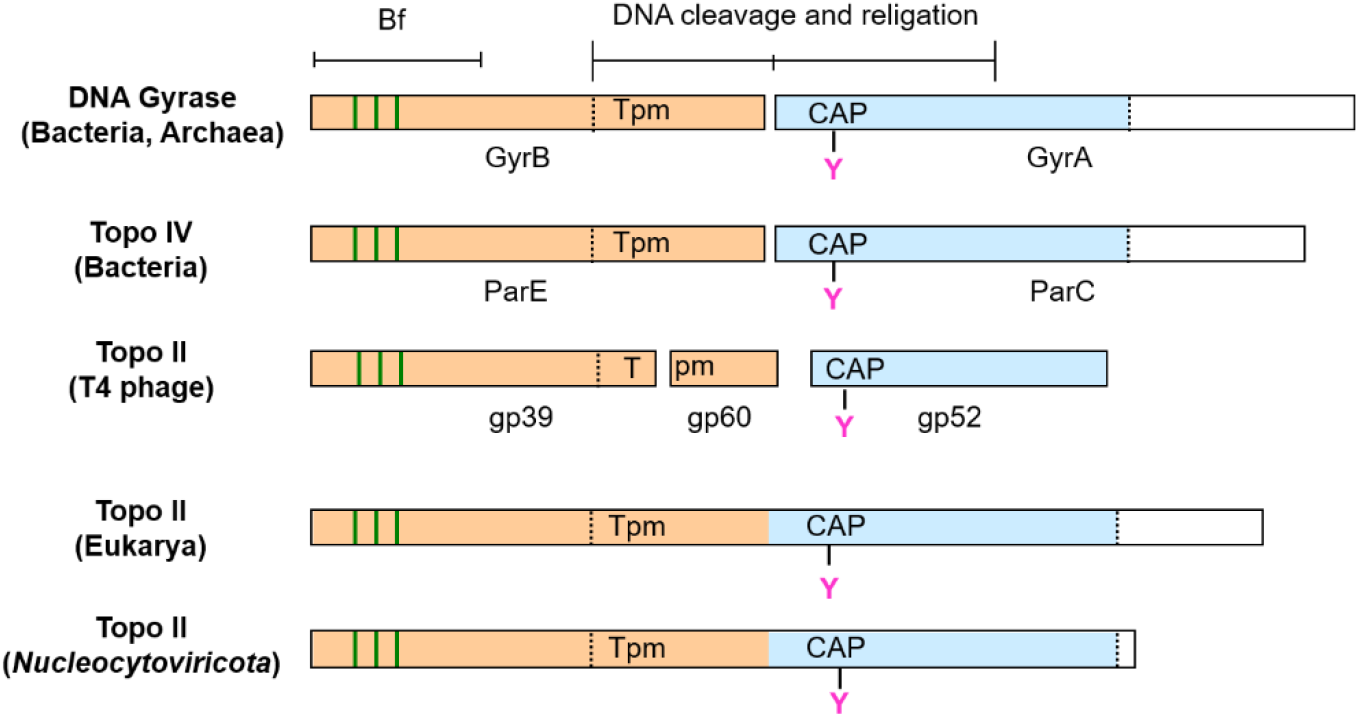
Schematic representation of the domain composition of Type II DNA topoisomerases of the family A. The different domains correspond to the Bergerat fold/ GHKL (Bf), TOPRIM (Tpm), the 5Y-CAP or winged helix (WHD) domain containing the catalytic Tyrosine.

A possible answer to this enigma could reside in the virosphere. The first viral Topo IIA was discovered in 1980 in the T4 bacteriovirus (Liu et al. 1979), the iconic virus of the class *Caudoviricetes* from the recently proposed family *Straboviridae*. Surprisingly, the T4 Topo IIA, a heterotrimer (Fig. 1), is not specifically related to bacterial Topo IIA, but branched between DNA gyrases and eukaryal Topo IIA in phylogenetic trees (Gadelle et al. 2003). Later on, Topo IIA genes were discovered in several members of the phylum *Nucleocytoviricota* (Fig. 1), formerly known as nucleocytoplasmic large DNA viruses (NCLDV)(Lavrukhin et al. 2000; Gadelle et al. 2003; Coelho et al. 2015; Coelho et al. 2016; Erives 2017). Inhibition of this Topo IIA disrupts replication of the African swine fever virus (ASFV; family *Asfarviridae*) *in vitro* (Freitas et al. 2016), indicating that compounds active against the ASFV-Topo IIA, such as fluoroquinolones, are promising drugs against the highly contagious and fatal disease caused in pigs by ASFV.

The Topo IIA encoded by *Nucleocytoviricota* are very similar to the ubiquitous eukaryotic Topo IIA at the sequence level and in that they are homodimers devoid of gyrase activity (Fig. 1). In the traditional view that considers viruses as pickpockets of cellular proteins, this suggests that Topo IIA were acquired by *Nucleocytoviricota* from their eukaryotic hosts. However, in the framework of the “out of viruses” hypothesis for the origin of DNA (Forterre 2002), it is tempting to suggest that this gene transfer took place the other way around, and that eukaryotic Topo IIA was acquired from the *Nucleocytoviricota* (Forterre and Gadelle 2009). A preliminary phylogenetic analyses provided ambiguous results: some Topo IIA from *Nucleocytoviricota* branched between T4 and Eukarya, suggesting that Topo IIA was indeed transferred from viruses to cells, whereas other viral enzymes branched within eukaryotes in agreement with transfers from cells to viruses (Gadelle et al. 2003; Forterre et al. 2007).

At the time of these analyses, only six Topo IIA from four families (*Asfarviridae, Mimiviridae, Iridoviridae, Phycodnaviridae*) within *Nucleocytoviricota* were known (Forterre et al. 2007). During the last decade, a great number of new *Nucleocytoviricota* genomes became available, including those of giant viruses from the families *Mimiviridae, Marseilleviridae* and *Pandoraviridae*, which encode Topo IIA (Abergel et al. 2015; Colson et al. 2017). Notably, it was shown that the Topo IIA encoded by *Marseilleviridae* branch as a sister clade to Eukarya (Erives 2017). We thus decided to update the Topo IIA phylogenetic classification, focusing on viral and eukaryotic Topo IIA. Our results strongly suggest that eukaryotic Topo IIA originated from a Topo IIA ancestor encoded by a virus closely related to modern *Megaviricetes*, a class of *Nucleocytoviricota* that includes many giant viruses, such as *Mimiviridae*. We have previously reported phylogenetic analyses suggesting that eukaryotic RNA polymerase II was probably recruited by eukaryotes from a virus related to *Imitervirales* in a tree including *Nucleocytoviricota* and the three nuclear RNA polymerases present in all eukaryotes (Guglielmini et al. 2019). One can speculate that both RNA polymerase II and Topo IIA were possibly acquired simultaneously by a proto-eukaryote in the lineage leading to the last eukaryotic common ancestor (LECA), in agreement with the fact that these two enzymes interact functionally and physically in modern eukaryotes. Regardless, our results support the hypothesis that interactions between proto-eukaryotes and *Nucleocytoviricota* have played an important role in shaping the physiology of modern eukaryotic cells.

## Material and methods

### Data collection

For Bacteria, we downloaded the full proteomes of a set of 112 bacterial strains spanning 10 representative groups (Aquificae, Dictyoglomi, Elusimicrobia, FCB group, Nitrospirae, PVC group, Proteobacteria, Spirochaetes, Terrabacteria group, Thermotogae). We used BLASTP v2.9.0+ (Ramsay et al. 2000; Camacho et al. 2009) recursively to collect the homologs of *Escherichia coli* K12 GyrB and GyrA proteins (WP_000072067.1, NP_416734.1) in those 112 proteomes. Finally, we concatenated each related pair of GyrB and GyrA hits. We also added the sequence of *E. coli* Topo IV (Table S1) that branched at the base of DNA gyrase in previous analyses.

For *Caudoviricetes* (tailed phages), we downloaded all the 1,131,926 *Caudoviricetes* proteins, and next used BLASTP to search for the homologs of the three subunits of the T4 phage topoisomerase II (E-value lower than 1e-10). We kept only *Caudoviricetes* lineages for which we obtained a hit for the three subunits, and concatenated the corresponding sequences. Interestingly, beside the group of previously known Topo IIA closely related to T4 Topo IIA infecting Enterobacteriaceae, we detected several new related Topo IIAs sequences in distantly related members of the *Myoviridae* infecting Rhizobiaceae and Firmicutes, but also in three members of the *Ackermannviridae* infecting Enterobacter and Rhizobiaceae as well as three unclassified siphoviruses infecting Firmicutes (Table S1).

For *Nucleocytoviricota* (NCLDV), we searched for Topo IIA sequences in *Nucleocytoviricota* genomes that we previously used to determine the list of core genes of *Nucleocytoviricota* and the phylogeny based on the concatenation of 8 core genes (Guglielmini et al. 2019). Topo IIA turned out to be present in all *Imitervirales* (formerly *Megavirales* in (Guglielmini et al. 2019)) an order that includes *Mimiviridae*, the so-called ‘extended Mimiviridae’ and several more recently described large DNA viruses (Catovirus, Hokovirus, Indivirus, Klosneuvirus and Tupanvirus) (Guglielmini et al. 2019). Topo IIA is also present in some members of the *Pimascovirales*, in particular, in all members of *Marseilleviridae* and viruses of the Pitho-like group (Orpheovirus, Cedratvirus, Pithovirus). Finally, Topo IIA is present in all members of the order *Asfuvirales*, which includes *Asfarviridae* and related viruses (Kaumovirus, Faustovirus, Pacmanvirus).

It turned out to be more challenging to assemble the dataset of eukaryotic Topo IIA sequences, because we found many fragments of Topo IIA in eukaryotic proteomes. Thus, we produced position-specific scoring matrices (PSSM) for GyrA and GyrB proteins using alignments from the PFAM database (files provided as supplementary materials) and searched for coding sequences matching both profiles. We defined a list of 52 eukaryotic organisms, representative of the known eukaryotic diversity. When possible, we downloaded the corresponding proteomes and used PSSM as PSIBLAST queries to obtain Topo IIA homologs. For those organisms where no proteome was available, we looked for transcriptomic data and performed *de novo* assembly using Trimmomatic v0.36 (Bolger et al. 2014) for the read pre-processing step, SortMeRNA v2.1b (Kopylova et al. 2012) to filter out rRNA sequences, and Trinity v2.2.0 (Grabherr et al. 2011) for the assembly. We then used the GyrA and GyrB PSSMs as queries for a TBLASTN search against the assemblies and kept hits matching both profiles.

### Phylogenetic analyses

All multiple amino-acid sequence alignments were performed using MAFFT v7.407 (Katoh and Standley 2013) and the E-INSi algorithm. Sites containing more than 50% of gaps were filtered out. Of note, for the tree with the largest taxonomic sampling, we used Divvier v1.0 (Ali et al. 2019) to reduce alignment errors with the MAFFT output.

All phylogenetic analyses were performed using IQ-TREE v1.6.7.2 (Nguyen et al. 2015). We selected the best-fit model using the IQ-TREE’s model finder (Wong et al. 2017) according to the BIC criterion. For the tree with the largest taxonomic sampling we used a mixture model (selected according to the BIC criterion) and the PMSF implementation (Wang et al. 2018). We made the search for the best tree more thorough by using the “allnni” option as well as setting the “pers” parameter to 0.2 and the “nstop” parameter 500. We always used 10 independent runs (--runs option of IQ-TREE) and selected the best one. Confidence branch supports were assessed using the transfer bootstrap expectation (1000 replicates except for the tree including all sequences, where 100 replicates were used (Lemoine et al. 2018). We used iToL v4.4.2 (Letunic and Bork 2016) to generate the figures. All trees and alignments are available https://doi.org/10.5281/zenodo.5702416.

## Results

### Viral Topo IIA branch between bacterial and eukaryotic Topo IIA in a global phylogeny

We first built a tree spanning the whole diversity of Topo IIA by including sequences from Bacteria, *Caudoviricetes, Nucleocytoviricota* and eukaryotic Topo IIA (Fig. 2). We did not include archaeal DNA gyrases because they branch within bacterial DNA gyrases in previous phylogenetic analyses (Forterre et al. 2007; Raymann et al. 2014). Importantly, we did not detect orthologues of eukaryotic-like Topo IIA in the MAGs of different lineages of Asgard archaea, but only bacterial-like DNA gyrases.

**Fig. 2.**
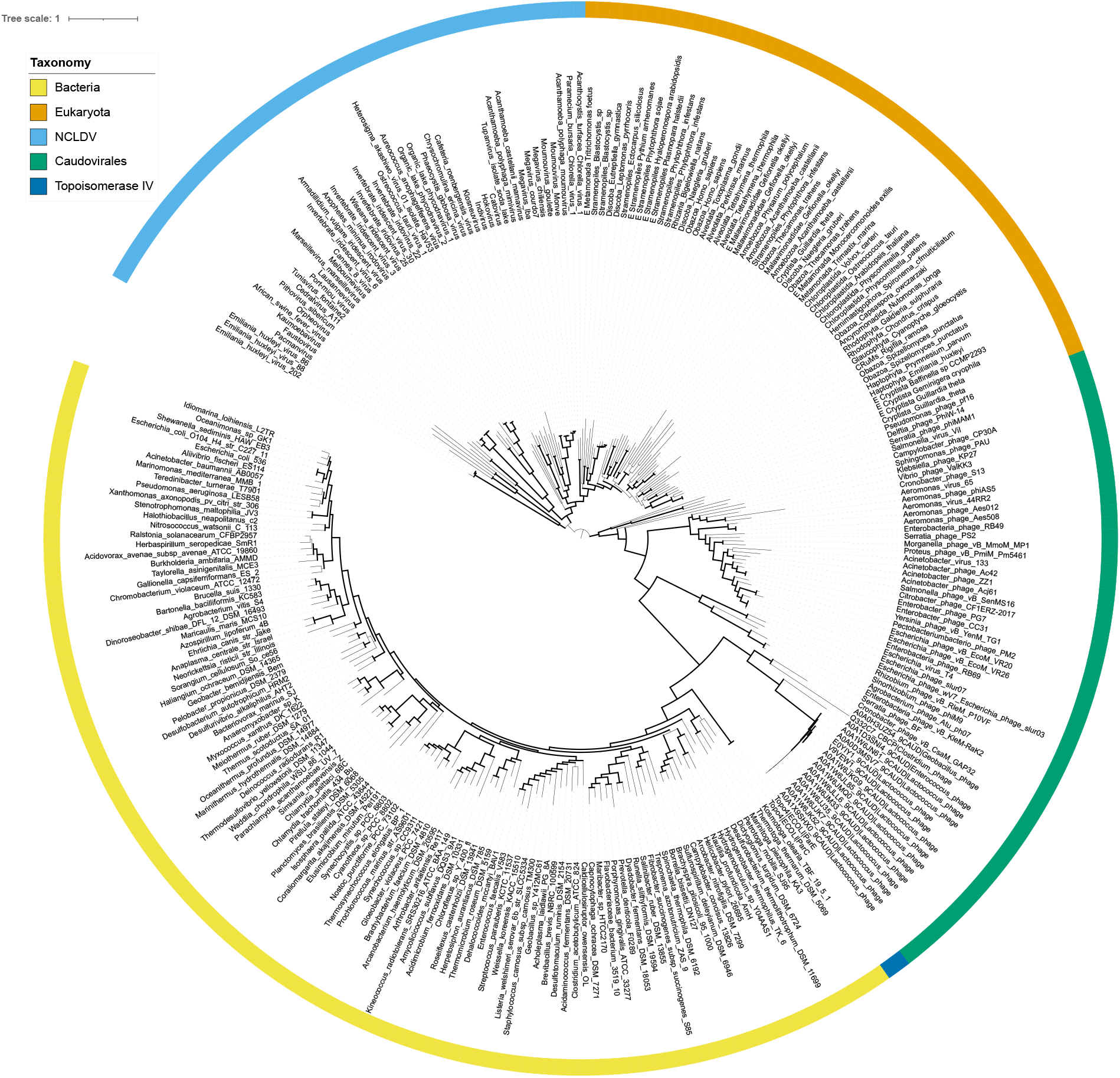
Phylogenetic tree of the Topo IIA. Maximum-likelihood tree for 269 Topo IIA proteins for bacteria (113 sequences, including the two Topoisomerase IV proteins from *Escherichia coli*, ParC and ParE), eukaryotes (53 sequences), *Nucleocytoviricota* (47 sequences) and *Caudoviricetes* (56 sequences). The outer circle colors represent the group to which the sequences belong. The selected model was LG+F+R15. Thick branches have a branch support (TBE) greater than 70%.

The four groups of sequences (Bacteria, *Caudoviricetes, Nucleocytoviricota* and eukaryotes) were clearly separated in the tree (Fig. 2). The tree was arbitrarily rooted between *Nucleocytoviricota* and *Caudoviricetes* for convenience, dividing the tree in two clusters, one grouping eukaryotes and their viruses (*Nucleocytoviricota)* and the other grouping Bacteria and their viruses (*Caudoviricetes)*. Both Bacteria and eukaryotes were monophyletic. In contrast, it was not possible to obtain the monophyly of either *Caudoviricetes* or *Nucleocytoviricota*. Importantly, in contrast to our previous analysis (Forterre et al. 2007), *Nucleocytoviricota* and eukaryotic Topo IIAs were not intermixed.

Although DNA viruses encode many viral-specific DNA replication proteins, they can sometimes recruit cellular replisome components (Krupovič et al. 2010). We thus wondered if the grouping of *Caudoviricetes* and *Nucleocytoviricota* between Bacteria and eukaryotes was due to the long-branch attraction (LBA) artifact, with *Caudoviricetes* branching within Bacteria and *Nucleocytoviricota* within eukaryotes. This seemed unlikely considering the great divergence between viral Topo IIAs and their cellular counterparts. However, to test this hypothesis, we built several subtrees, both to remove groups with long branches and to enhance the signal by increasing the number of meaningfully aligned amino acids. After removing bacterial sequences, the most distant outgroup, we obtained a tree topology largely reproducing the relationships between *Caudoviricetes, Nucleocytoviricota* and eukaryotes observed in the global phylogeny (Fig. 3). More importantly, after removing both Bacteria and *Caudoviricetes, Nucleocytoviricota* remained well separated from eukaryotes (Fig. 4 a, b). This indicates that the separation of *Nucleocytoviricota* and eukaryotes in the tree was not due to an attraction of *Nucleocytoviricota* by *Caudoviricetes* and/or Bacteria. Similarly, *Caudoviricetes* remained well separated from bacteria after removing both *Nucleocytoviricota* and eukaryotes (Fig. 5).

**Fig. 3.**
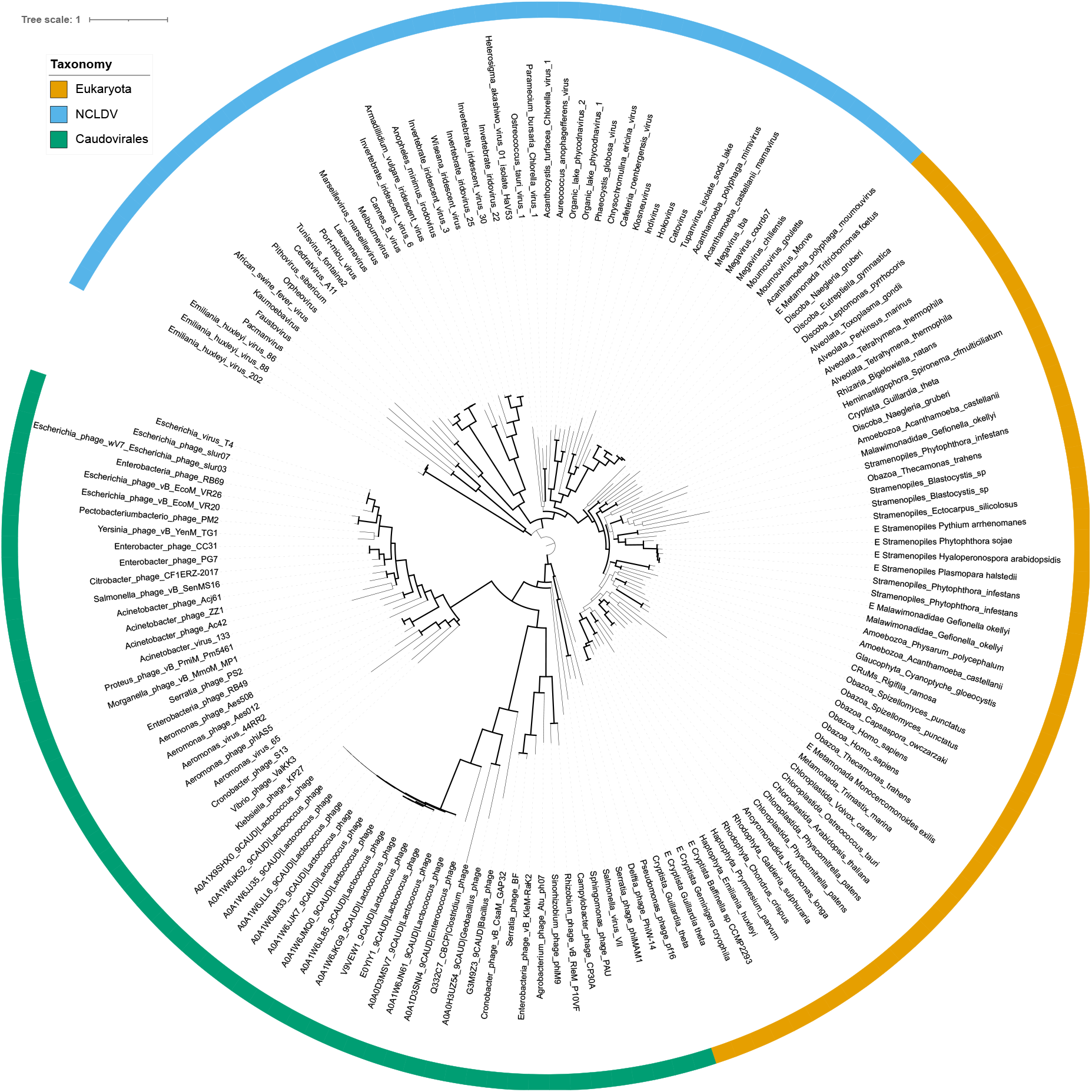
Phylogenetic tree of the Topo IIA for the eukaryotes, *Nucleocytoviricota* and *Caudoviricetes*. Maximum-likelihood tree for 156 Topo IIA proteins for eukaryotes (53 sequences), *Nucleocytoviricota* (47 sequences) and *Caudoviricetes* (56 sequences). The outer circle colors represent the group to which the sequences belong. The selected model was LG+F+R10. Thick branches have a branch support (TBE) greater than 70%.

**Fig. 4.**
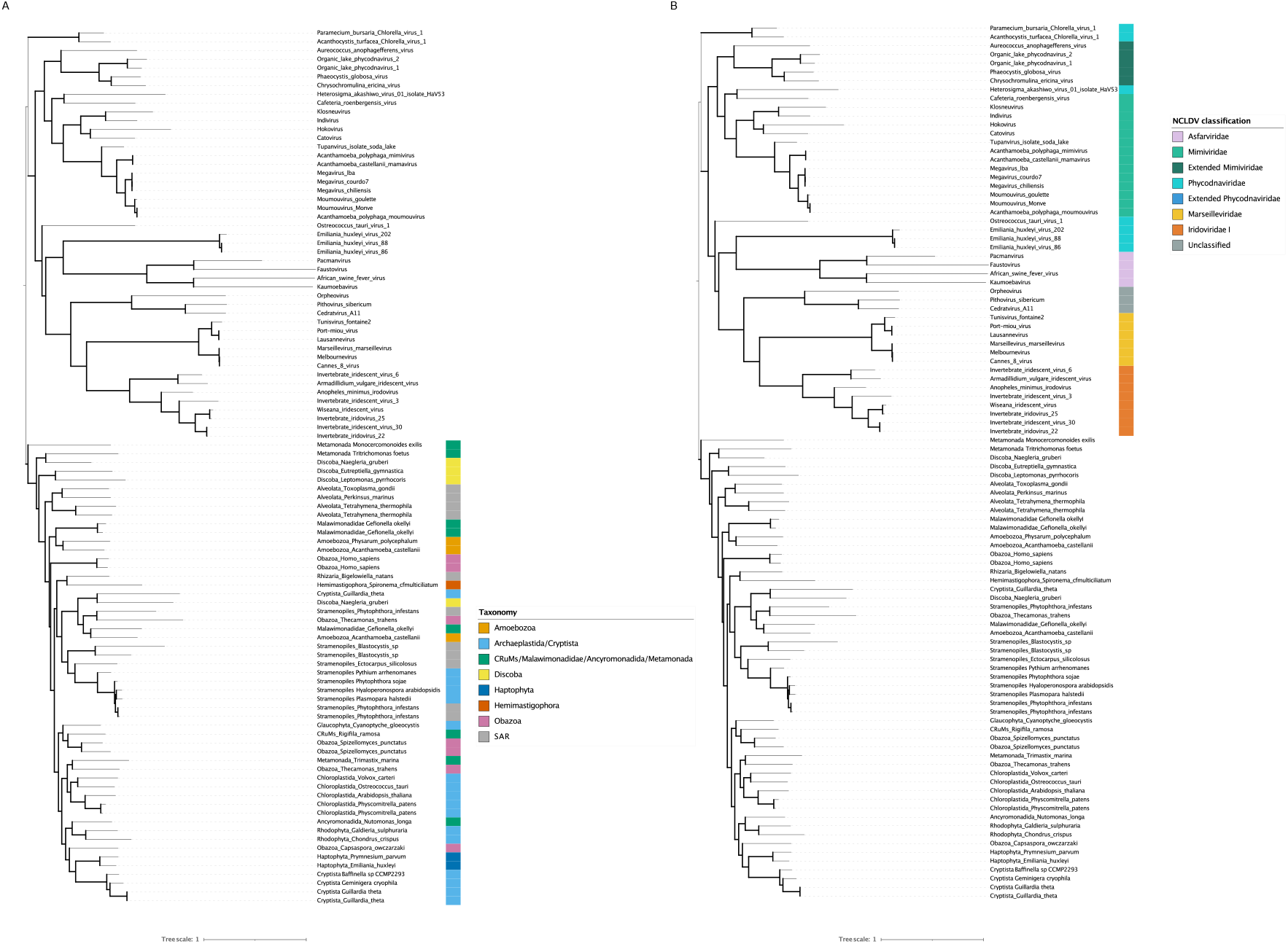
Phylogenetic tree of the Topo IIA for the eukaryotes and *Nucleocytoviricota*. Maximum-likelihood tree for 100 Topo IIA proteins for eukaryotes (53 sequences) and *Nucleocytoviricota* (47 sequences). The selected model was LG+F+R6. Thick branches have a branch support (TBE) greater than 70%. In panel A, the colored bar represents the eukaryotes classification. In panel B, the colored bar represents the *Nucleocytoviricota* classification.

**Fig. 5.**
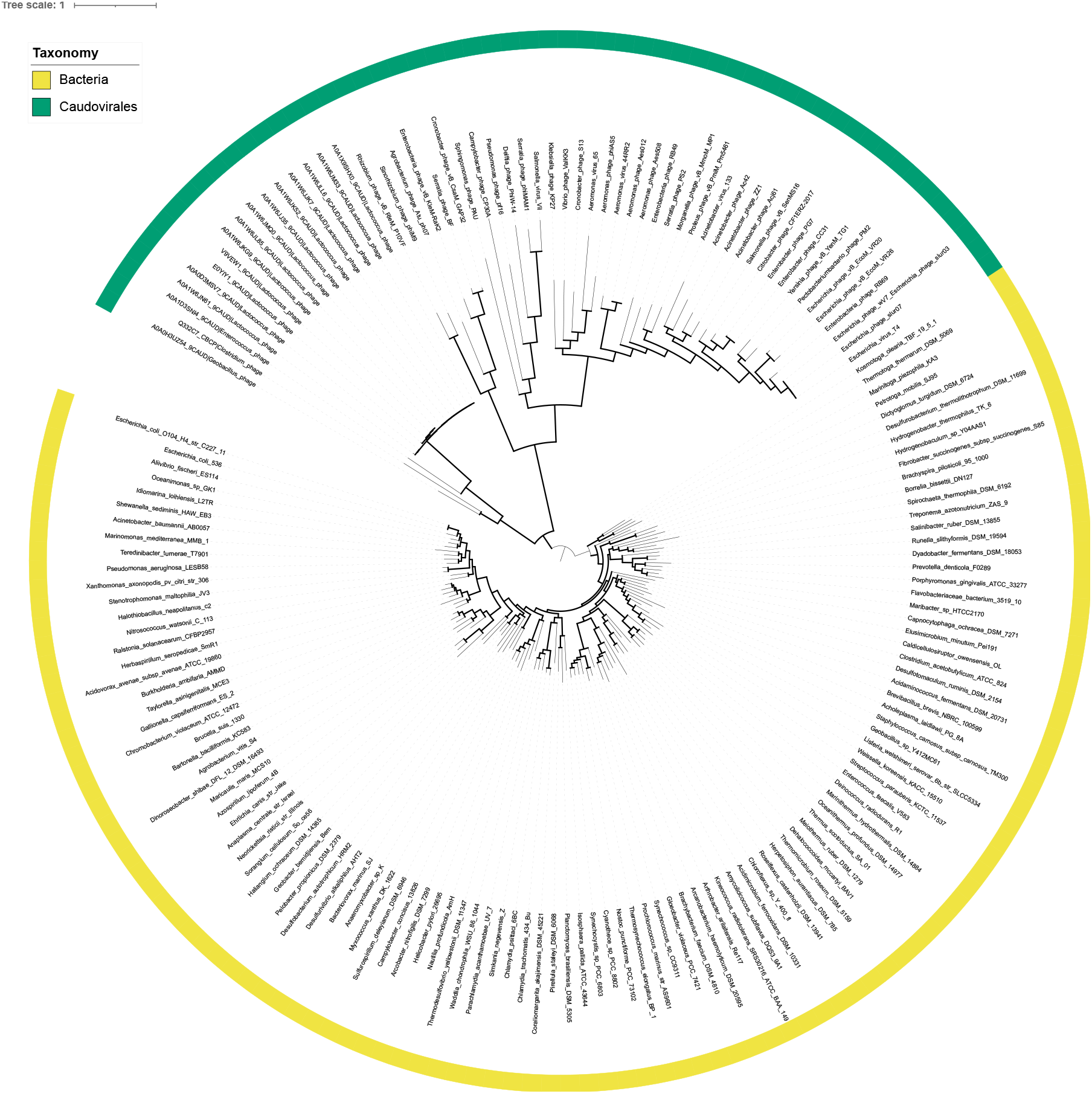
Phylogenetic tree of the Topo IIA for the bacteria and *Caudoviricetes*. Maximum-likelihood tree for 167 Topo IIA proteins for bacteria (111 sequences) and *Caudoviricetes* (56 sequences). The outer circle colors represent the group to which the sequences belong. The selected model was LG+R11. Thick branches have a branch support (TBE) greater than 70%.

The distribution and phylogeny of Topo IIA can provide information about their presence, or not, in the ancestors of each group of organisms. The ubiquity of DNA gyrase in bacteria leaves little doubt that this enzyme was present in the Last Bacterial Common Ancestor (LBCA). Similarly, the ubiquity of the single-subunit Topo IIA in Eukarya testifies to the presence of at least one Topo IIA in the LECA. However, we did not recover the monophyly of all major eukaryotic divisions in our phylogeny (Fig. 4a). Members of certain divisions were present in different parts of the tree, suggesting a complex history of Topo IIA during the diversification of eukaryotes, including gene duplication and gene loss. Several eukaryotes indeed contain more than one Topo IIA gene (Forterre et al. 2007). Some correspond to recent duplications (such as the Topo IIα and Topo IIβ in vertebrates), but others probably correspond to more ancient gene duplications or possibly gene transfers between eukaryotic lineages. With the root of the eukaryotic tree being still debated (Burki et al. 2019), it is difficult to propose a scenario for the evolution of Topo IIAs in eukaryotes. From our phylogenetic analyses, one cannot exclude that LECA already contained more than one Topo IIA.

The broad representation of Topo IIA in *Nucleocytoviricota* suggests that this enzyme was also present in the Last *Nucleocytoviricota* Common Ancestor (LNCA), and was subsequently lost in a few lineages. This hypothesis is supported by the congruence between the phylogenetic tree of *Nucleocytoviricota* Topo IIA (Fig. 4b) and the global phylogenetic classification of *Nucleocytoviricota* based on the concatenation of eight (core) genes present in most families of this phylum (Guglielmini et al. 2019). In the 8-core-genes phylogeny, *Nucleocytoviricota* were divided into two clusters that we named PAM (*Phycodnaviridae, Asfarviridae, Mimiviridae)* and MAPI (*Marseilleviridae, Ascoviridae*, Pitho-like viruses, *Iridoviridae)*, respectively. The PAM cluster included viruses corresponding to the recently proposed class *Megaviricetes* and *Pokkesviricetes*, whereas the MAPI cluster corresponded to the recently proposed order eukaryotes (Fig. 4b), we recovered the monophyly of all families of *Nucleocytoviricota*, as well as and the monophyly of *Pimascovirales*. Notably, the NCLDV Topo IIA tree was rooted deep within *Megaviricetes* when eukaryotes were used as the outgroup.

In contrast to the situation with Bacteria, eukaryotes and *Nucleocytoviricota*, Topo IIA are only present in a few subgroups of *Caudoviricetes*. Most Topo IIA are encoded by T4-like myoviruses (i.e., viruses with contractile tails recently reclassified into the family *Straboviridae*) with larger genomes, suggesting that Topo IIA was present in the last common ancestor of this phage group. Topo IIA encoded by *Ackermannviridae* (anther family of phages with contractile tails) branched within *Straboviridae* suggesting lateral gene transfer between these viral families (Fig. 5b). Three of the four Topo IIA encoded by viruses infecting Firmicutes have been tentatively assigned to the family *Siphoviridae* (phages with long non-contractile tails). They were grouped with Topo IIA of T4-like viruses, as a sister clade of bacterial homologs if the tree is rooted between *Nucleocytoviricota* and *Caudoviricetes*.

## Discussion

To discuss possible evolutionary scenarios, we arbitrarily rooted the Topo IIA phylogenetic tree (Fig. 2) at the three possible positions between the four clusters (Fig. 6 a,b,c). Rooting the tree between *Nucleocytoviricota* and eukaryotes (Fig. 6a) would suggest that *Nucleocytoviricota* and eukaryotic Topo IIA originated from a common viral or cellular ancestor. This scenario appears unlikely since it also implies that *Caudoviricetes* Topo IIA originated from *Nucleocytoviricota* Topo IIAs and *in fine* that bacterial DNA gyrases themselves originated from *Nucleocytoviricota* via *Caudoviricetes*. In that case, one should imagine that the LBCA originated after the diversification of *Nucleocytoviricota*. Since this diversification took place before LECA, at the time when ancestral *Nucleocytoviricota* infected proto-eukaryotic hosts, this scenario would suggest that proto-eukaryotes evolved before bacteria.

**Fig. 6.**
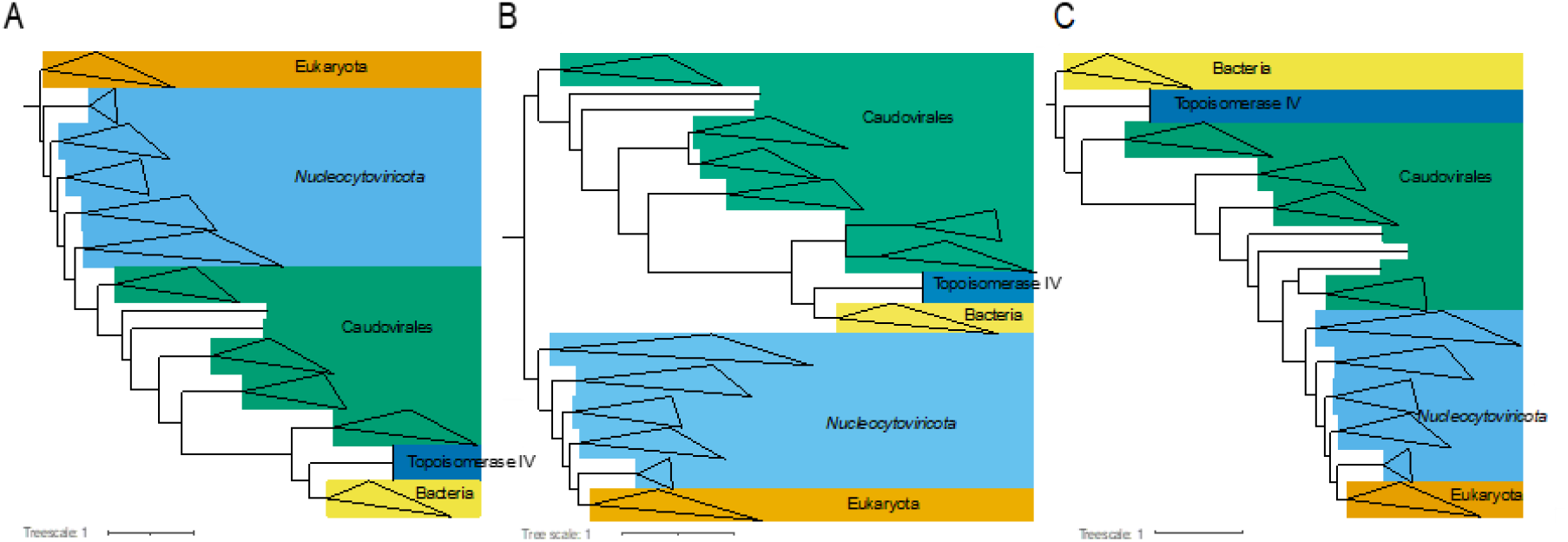
Schematic representation of the different possible rooting for the Topo IIA phylogenetic tree. The tree is the same as in Fig. 2. Monophyletic clades encompassing sequences from the same group have been collapsed and colored following Fig. 2. Panel A: rooting using the eukaryotes as an outgroup. Panel B: rooting using the bacteria and *Caudoviricetes* as an outgroup. Panel C: rooting using the bacteria as an outgroup.

Rooting the tree between *Caudoviricetes* and *Nucleocytoviricota* (Fig. 6b) produced two clusters corresponding to Bacteria/*Caudoviricetes* and to *Nucleocytoviricota*/eukaryotes. This rooting suggests that bacterial DNA gyrase originated from Topo IIA of *Caudoviricetes*, whereas eukaryotic Topo IIA originated from those of *Nucleocytoviricota*. Considering the universal conservation of Topo IIA in Bacteria and eukaryotes, this scenario suggests that the transfer from viruses to cells took place before the emergence of the LBCA and LECA, respectively. Hence, both *Caudoviricetes* and *Nucleocytoviricota* should have originated and diversified before the emergence of the LBCA and LECA, infecting proto-bacterial and proto-eukaryotic hosts, respectively. Such ancient origin would explain the great divergence between the different versions of Topo IIA. The diversification of *Nucleocytoviricota* before LECA is indeed supported by the RNA polymerase phylogenetic tree including both viral and cellular enzymes (Guglielmini et al. 2019). Moreover it has been suggested that *Caudoviricetes*, which also infect archaeal hosts (Liu et al. 2021), have diverged even prior to the emergence of LUCA (Krupovic et al. 2020).

Rooting the tree between Bacteria and *Caudoviricetes* (Fig. 6c) produced a tree in which Topo IIA of Bacteria and *Caudoviricetes* diverged from a common ancestor that predated the LBCA. In that case, the *Caudoviricetes* Topo IIA would have diverged from their bacterial counterparts before LBCA and continued during the diversification of Bacteria. The tree of Fig. 6c is consistent with the scenario in which eukaryotic viruses originated from a melting pot of bacterial viruses that infected the bacterium at the origin of mitochondria (Koonin, Krupovic, et al. 2015; Koonin, Dolja, et al. 2015) or another ancient bacterial endosymbiont present in a proto-eukaryotic ancestor of modern eukaryotes. In that case, the Topo IIA from a *Caudoviricetes* present in this putative bacterial endosymbiont would have been transferred to an ancestor of *Nucleocytoviricota*, potentially with other components of the DNA replication machinery shared between *Caudoviricetes* and *Nucleocytoviricota*, including NAD-dependent DNA ligase (Yutin and Koonin 2009). Notably, comparison of DNA replication machineries of all dsDNA viruses revealed a strong evolutionary and likely functional coupling between DNA topoisomerases and DNA ligases, with 96% of viruses encoding DNA topoisomerases also carrying a gene for a ligase (Kazlauskas et al. 2016). To explain the great divergence between the Topo IIA encoded by *Caudoviricetes* and those encoded by *Nucleocytoviricota* in terms of sequences and structure, this scenario entails that the rate of Topo IIA evolution increased dramatically following the transfer of the *Caudoviricetes* version into the lineage leading to the LNCA, with the fusion of the three Topo IIA subunits of *Caudoviricetes* Topo IIA into a single polypeptide.

The trees of Fig. 6b and 6c both can be also interpreted in the framework of the “*out of virus hypothesis*” for the origin of DNA topoisomerases (Forterre 2002; Forterre and Gadelle 2009). Conceivably, the different versions of Topo IIA originated in an ancient viral world. The scenario illustrated in Fig. 6b explicitly posits that proto-bacteria acquired their Topo IIA from an ancient *Caudoviricetes*, whereas in Fig. 6c, the bacterial and *Caudoviricetes* Topo IIA evolved from a common ancestor, which may or may not have been a virus. Regardless, in both scenarios, the eukaryotic Topo IIA has been acquired from *Nucleocytoviricota*. We have previously proposed a similar scenario to explain the ubiquitous distribution of Topo VI in Archaea and the restricted distribution of Topo VIII (both members of the Topo IIB family) in some archaea and bacterial mobile elements (Gadelle et al. 2014). In that scenario, the restricted distribution of Topo IIA to a few subgroups of *Caudoviricetes* seems surprising, but it resembles the restricted distribution of a recently described new version of RNA polymerase in a subgroup of these viruses (Weinheimer and Aylward 2020).

Importantly, if we exclude the unlikely conjecture in which the Topo IIA phylogenetic tree is rooted between *Nucleocytoviricota* and eukaryotes (Fig. 6a), the branching of all eukaryotes within *Nucleocytoviricota* in all other configurations suggests that a Topo IIA was introduced into eukaryotes from a member of this viral phylum. If the node at the base of the eukaryotic monophyletic clade corresponds to the position of LECA, as expected from the ubiquity of this enzyme in eukaryotes, the transfer of Topo IIA should have occurred before the emergence of LECA, i.e., from a member of *Nucleocytoviricota* to a proto-eukaryote. Alternatively, Topo IIA could have been introduced from *Nucleocytoviricota* to a particular eukaryotic lineage and later transferred from this lineage to all other lineages by horizontal gene transfer. This last scenario seems unlikely considering that Topo IIAs are present in all contemporary lineages of eukaryotes, without exception, and the enzyme is essential for several key functions conserved in all eukaryotic lineages. The fact that eukaryotes emerge in our analysis within the PAM group is consistent with the possibility that divergence of *Nucleocytoviricota* into several major families has predated the emergence of LECA (Guglielmini et al. 2019).

Our result raises an interesting question: which Topo II did proto-eukaryotes use before they captured the viral Topo IIA? A likely answer is that they relied on Topo IIB, since this enzyme is ubiquitous in Archaea, but also present in many eukaryotes. A Topo IIB-like protein with a very divergent V-B subunit is present in all eukaryotes and is part of the complex responsible for initiation of meiotic recombination (Vrielynck et al. 2016; Robert et al. 2017) whereas several eukaryotic lineages, e.g., Viridiplantae, contain a *bona fide* archaeal-like Topo IIB (Forterre et al. 2007; Malik et al. 2007; Forterre and Gadelle 2009).

In comparing the *Nucleocytoviricota* core genes’ phylogeny with the phylogeny of the three eukaryotic nuclear RNA polymerases and those of *Nucleocytoviricota*, we have previously shown that two of the eukaryotic RNA polymerases, Pol I and Pol II, were probably introduced into the proto-eukaryotic lineage from *Nucleocytoviricota* (Guglielmini et al. 2019). This possibility was strongly supported in the case of the RNA polymerase II. It is worth noting that, like the position of Topo IIA in the present study, the RNA Pol II branched within *Megaviricetes* in the RNA polymerase tree. One can speculate that these two proteins (that play a major role in the eukaryotic transcription machinery) were recruited together from the same virus. This would make sense from the viewpoint of cell physiology, since the two enzymes interact both functionally and structurally. Indeed, it has been shown that Topo IIA is a structural component of the holo-Pol II complex and is essential for efficient RNA synthesis of nucleosomal DNA by this complex (Mondal and Parvin 2001). Topo IIA is required to produce long RNA Pol II transcripts (>3 kb) in *Saccharomyces cerevisiae* (Joshi et al. 2012) and enhances the recruitment of RNA Pol II to promoters in budding yeast (Sperling et al. 2011). It is possible that both Topo IIA and RNA Pol II were domesticated by a proto-eukaryote following the integration of a *Nucleocytoviricota* encoding these genes into the host chromosome. Integration of entire or large portions of the genomes of some *Nucleocytoviricota* into the chromosome of modern eukaryotes has been well documented (Delaroque and Boland 2008; Cock et al. 2010; Filée 2014; Moniruzzaman et al. 2020).

The viral origin of eukaryotic Topo IIA, in addition to those of RNA Pol II and possibly RNA Pol I, strengthens the idea that giant viruses of the phylum *Nucleocytoviricota* (especially members of the PAM group) played a major role in shaping the identity of modern eukaryotes (Forterre and Gaïa 2016). It is likely that other important proteins involved in eukaryotic physiology originated from *Nucleocytoviricota*. This has been proposed for eukaryotic histones, since the four histones from Medusavirus and Marseilleviruses branch at the base of the eukaryotic clades of their respective homologues (Erives 2017; Yoshikawa et al. 2019) and for enzymes involved in mRNA capping (Bell 2020). However, in those cases, robust phylogenetic analyses remain to be carried out since the published papers are based on limited sampling of the eukaryotic and *Nucleocytoviricota* diversity. The viral origin of some of the major players in eukaryotic cell biology was probably not limited to nuclear components since we have recently reported that the eukaryotic cytoplasmic actin might have been recruited by proto-eukaryotes from an actin-like protein (viractin) encoded by some *Imitervirales*, an order of *Megaviricetes* (Da Cunha et al. 2020).

The eukaryotic molecular fabric appears to be a melting pot of proteins that originated in *Nucleocytoviricota* (mainly *Megaviricetes*), those that emerged *de novo* in the eukaryotic stem branch, proteins inherited from the bacterial ancestor of mitochondria and chloroplasts, and proteins that had ancestors in Archaea (in two domains scenarios) or in the common ancestor of Archaea and eukaryotes (in three domains scenario). Sorting out the viral component of our eukaryotic ancestors is now a major task in understanding eukaryogenesis.

## Supporting information

GyrA-GyrB PSSM

## Author contributions

Conceptualization: JG, MG, VDC, PF

Methodology: JG, AC, PF

Investigation: JG, MG, PF

Supervision: AC, MK, PF

Writing – original draft: JG, PF

Writing – review & editing: JG, MG, VDC, AC, MK, PF

## Data and materials availability

All data are available in the main text or upon request.

